# CRISPR-Cas9 genome editing in *Steinernema* entomopathogenic nematodes

**DOI:** 10.1101/2023.11.24.568619

**Authors:** Mengyi Cao

## Abstract

Molecular tool development in traditionally non-tractable animals opens new avenues to study gene functions in the relevant ecological context. Entomopathogenic nematodes (EPN) *Steinernema* and their symbiotic bacteria of *Xenorhabdus* spp are a valuable experimental system in the laboratory and are applicable in the field to promote agricultural productivity. The infective juvenile (IJ) stage of the nematode packages mutualistic symbiotic bacteria in the intestinal pocket and invades insects that are agricultural pests. The lack of consistent and heritable genetics tools in EPN targeted mutagenesis severely restricted the study of molecular mechanisms underlying both parasitic and mutualistic interactions. Here, I report a protocol for CRISPR-Cas9 based genome-editing that is successful in two EPN species, *S. carpocapsae* and *S. hermaphroditum*. I adapted a gonadal microinjection technique in *S. carpocapsae*, which created on-target modifications of a homologue *Sc-dpy-10* (cuticular collagen) by homology-directed repair. A similar delivery approach was used to introduce various alleles in *S. hermaphroditum* including *Sh-dpy-10* and *Sh-unc-22* (a muscle gene), resulting in visible and heritable phenotypes of dumpy and twitching, respectively. Using conditionally dominant alleles of *Sh-unc-22* as a co-CRISPR marker, I successfully modified a second locus encoding Sh-Daf-22 (a homologue of human sterol carrier protein SCPx), predicted to function as a core enzyme in the biosynthesis of nematode pheromone that is required for IJ development. As a proof of concept, *Sh-daf-22* null mutant showed IJ developmental defects *in vivo* (*in insecta)*. This research demonstrates that *Steinernema* spp are highly tractable for targeted mutagenesis and has great potential in the study of gene functions under controlled laboratory conditions within the relevant context of its ecological niche.

## Introduction

Genetic models shaped the foundation of our understanding of biology. Classical invertebrate model animals, such as *Caenorhabditis elegans* and *Drosophila melanogaster,* enabled well-controlled experiments to interrogate gene functions, which revealed conserved signaling pathways among diverse species including humans. However, the complete understanding of eukaryotic gene functions is reliant on examining phenotypes within the animal’s natural context, such as its native biosphere, a specific host animal environment, or the presence of natural microbiota (Collins et al., 2021; Ebert & Fields, 2020; McFall-Ngai et al., 2013). Such ecological relevance is difficult to be fully addressed in most of the classical genetic model animals because they were particularly chosen to simplify the complicated natural context using laboratory-specific conditions (Petersen et al., 2015). Therefore, it is crucial to establish genetic tools in non-traditional model animals, in which novel functions of unique and conserved genes could be studied in the relevance of ecological niche, and maintainable in a laboratory setting for controlled experimentation.

Genetic tool development is particularly valuable in the study of animal-microbes symbiosis. In the past decades, the establishment of molecular genetics in diverse bacterial species expanded our knowledge in interactions among host animals and microbes, enabling the study of microbial physiology in host niches at the level of organs and tissues (Ruby, 2008). Recently, functional genomic tools are successfully adapted to animals that are traditionally non-tractable and revolutionized the study of evolutionary biology and ecology (Gudmunds et al., 2022). The establishment of these molecular tools in diverse animal species facilitated the investigation of gene functions in response to the ecological context, including the signals that they receive from their host animal and native symbionts (Atkinson & Hallem, 2022; Chung et al., 2018; Cleves et al., 2020; Klimovich et al., 2019).

The soil-dwelling and entomopathogenic nematode (EPN) of *Steinernema* spp are suitable experimental systems to investigate animal physiology in the ecological context because they can be studied in the field, within the infected insect, and *in-vitro* on petri-dishes in the presence or absence of its native symbiotic bacteria. During the infective juvenile stage (IJ) stage, *Steinernema* nematode carries its mutualistic symbiotic bacteria in the intestinal pocket (receptacle) in a species-specific manner (one species of bacteria associates with one species of animal) and seeks for an insect host via a repertoire of host-seeking behaviors (Forst & Clarke, 2002; Goodrich-Blair & Clarke, 2007; Hallem et al., 2011; Murfin, Dillman, et al., 2012). Once invading an insect, IJ releases its symbiotic bacteria, and both partners kill the insect. *Xenorhabdus* bacteria replicate using the nutrient from the insect cadaver and the bacteria serve as food for the nematodes (Mucci et al., 2022), supporting their reproductive development (juveniles J1-J4 and adults). The depletion of nutrients and nematode overcrowding within the dead insect, causing accumulation of pheromone, could induce IJ development, and promote IJs disperse from the insect cadaver into the soil to seek for a new insect prey (Kaplan et al., 2012; Popiel et al., 1989) (Fig. 1A). In the laboratory, the *Steinernema* nematodes can be maintained through insects to recapitulate their natural life cycles (White, 1927). The nematode and bacterial symbiont can also be cultured separately and together, facilitating the study of host-microbe interactions (Mitani et al., 2004; Vivas & Goodrich-Blair, 2001). *Steinernema* nematodes are applicable for field studies: certain *Steinernema* spp are commercially available as non-pesticide control agents to antagonize insect pests and promote agricultural productivity (Ehlers, 2001; Shapiro-Ilan et al., 2012; Tarasco et al., 2017).

**Figure 1:**
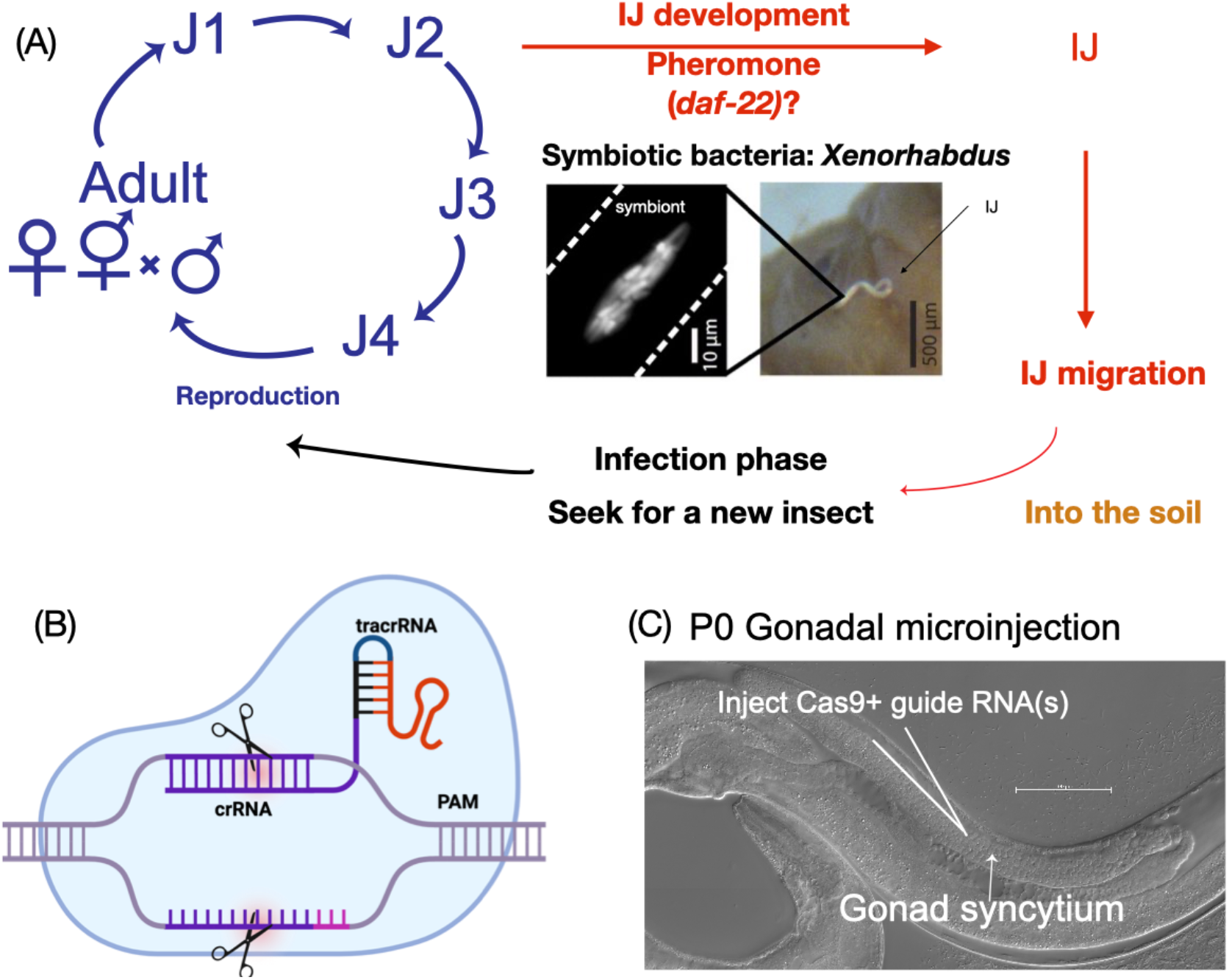
*Steinernema* nematodes life cycle and overview of CRISPR-Cas9 based genome-editing via gonadal microinjection. (A): Life cycle of *Steinernema* nematodes. Infective juveniles (IJs) are associated with symbiotic bacteria in the intestinal pocket (receptacle). They migrate in the soil and infect an insect. Within the infected insect, IJs release symbiotic bacteria, and both partners kill the insect. Nematodes reproduce inside an insect cadaver feeding on symbiotic bacteria. The reproductive cycle includes four juvenile stages J1-J4. Adults are either male-female or male-hermaphrodite. Overcrowding and accumulation of pheromone induces IJ development. ((B): Cas9 ribonucleoprotein (RNP) is a complex of guide RNA (sgRNA containing crRNA and tracrRNA) and Cas9 protein (blue bubble). crRNA matches 20 nucleotides upstream of the PAM site, guiding the cleavage on double-stranded DNA three base pairs upstream of PAM (adapted from “Cas9-sgRNA-cleavage”, by BioRender.com (2023)). (C): Microinjction of RNP into the gonad of *S. hermaphroditum*.

Genetic tools, such as transposon-based mutagenesis screen and gene knockouts, are currently available in multiple *Xenorhabdus* bacterial species, and have been used to study symbiont gene functions in the context of the parasitic and mutualistic host animals (Alani et al., 2023; Herbert & Goodrich-Blair, 2007; Murfin et al., 2019; Vivas & Goodrich-Blair, 2001). Our recent work in an isolate of *Steinernema hermaphroditum* (PS9179) established a convenient breeding and efficient cryopreservation protocol and a plausible forward genetics screen pipeline, which showcases the genetic tractability of the animal host (Cao et al., 2021). However, in EPNs including *Steinernema*, there are no consistent and heritable molecular tools for targeted mutagenesis (reverse genetics), which hampered the in-depth study of gene functions on the nematode side.

The clustered regularly interspaced short palindromic repeats (CRISPR) and *Streptococcus pyogenes* Cas9 system allowed precise genome-editing in traditionally non-tractable experimental systems (Fig. 1C) (Wiedenheft et al., 2012). Guided by RNA, the Cas9 endonuclease creates a double-stranded DNA break, allowing modification at targeted locus in the genome. A Cas9 protein assembles with two small RNAs, a CRISPR RNA (crRNA), and a *trans-*activating crRNA (tracrRNA), to create a functional Cas9 ribonucleoprotein (RNP). At the targeted site where a protospacer adjacent motif (PAM, 5’-NGG-3’) is present, crRNA matches and recognizes the twenty nucleotides upstream of the PAM site, leading to Cas9 cleavage three base pairs upstream of the PAM site. The double-stranded break could be repaired by error-prone non-homologous end joining (NHEJ) or precise homology-directed repair (HDR).

The CRISPR-Cas9 system has been successfully adopted in various nematode species, including the classical model *C. elegans,* the necromenic nematode *Pristionchus pacificus,* the mammalian parasite *Strongyloides stercoralis,* and other non-model species, such as *Aunema*, *Oscheius,* and *Panagrolaimus* (Adams et al., 2019; Castelletto & Hallem, 2021; Dockendorff et al., 2022; Hellekes et al., 2023; Hiraga et al., 2021; H.-M. Kim et al., 2022; Lok, 2019). In these successful cases, either plasmid encoded Cas9 and guide RNAs or Cas9 RNP were delivered through microinjection into the gonad of a hermaphrodite or a female nematode. The site of injection is the central core of the distal arm of the gonad, termed the gonadal syncytia, that contains a cytoplasm that is shared by many germ cell nuclei (Evans, 2006). This approach, adapted from *C. elegans* transgene and RNAi (C. C. Mello et al., 1991; C. Mello & Fire, 1995), is effective because injected reagents can be delivered to multiple progenies. This microinjection approach is also used in delivering double-stranded RNA to entomopathogenic nematode *Heterorhabditis* which showed success in RNAi of first-generation progeny (Ratnappan et al., 2016). Recently, co-CRISPR and co-conversion methods were adapted to the CRISPR-Cas9 genome-editing in nematodes to enrich genome alteration that does not necessarily cause an immediately visible phenotype. Using a co-CRISPR or co-conversion locus with a dominant and visible phenotype, the animals showing modification of the marker would be more likely to edit a second independent locus (Arribere et al., 2014; H. Kim et al., 2014).

In this research, I developed a CRISPR-Cas9 based approach to introduce on-target modifications in *Steinernema* nematodes. To deliver Cas9 RNP that is assembled *in vitro,* I adopted the gonadal microinjection techniques from established nematode genetic models. This method is successful in both *S. carpocapsae* and *S. hermaphroditum,* with evidence of HDR and NHEJ. Homozygous alleles of *dpy-10* and *unc-22* in *S. hermaphroditum* could be maintained *in-vitro* and showed expected phenotypes on petri-dishes. I used conditionally dominant allele *Sh-unc-22* as a co-CRISPR marker and successfully enriched the modification of another gene, *Sh-daf-22,* which showed expected IJ (dauer larvae) developmental defect within the infected insects. This study demonstrates *Steinernema* as a highly tractable model for reverse genetics, and it has great potential to study conserved gene functions in controlled laboratory conditions and relevant ecological context.

## Materials and Method

### Bacterial growth and nematode maintenance

Bacterium *Xenorhabdus griffiniae* (HGB2511), *Xenorhabdus nematophila* (HGB800), *E. coli* OP50 and *Comammonas acquatica* (DA1877) were grown on Luria Broth (LB) agar supplemented with 1% sodium pyruvate (Xu & Hurlbert, 1990). Single colonies of bacteria were used to start overnight cultures in dark LB media at 30°C with aeration. Bacterial overnight culture was used to seed patches on NGM (Brenner, 1974) and lipid agar (Vivas & Goodrich-Blair, 2001). *Steinernema hermaphroditum* wild-type and mutant lines were maintained at 22-25°C feeding bacteria (either *Commamonas* or *Xenorhabuds griffiniae*) grown on NGM agar plates. *Steinernema carpocapsae* (All strain) infective juveniles (IJs) were maintained through *Galleria mellonella* 5^th^ instar insects (PetSmart, Phoenix, AZ) by white-trapping (White, 1927). Conventional IJs were surface-sterilized in 1% bleach, washed with sterile water for three times, and recovered on *Xenorhabduds nematophila* bacterial lawn grown on lipid agar plates at 25°C.

### Target gene sequence analysis and Cas9 ribonucleoprotein preparation

The sequence of *Sc-dpy-10* was extracted from *S. carpocapsae* complete genome (BioProject PRJNA202318) using WormBase ParaSite (Howe et al., 2016, 2017). Clustal Omega Multiple Sequence Alignment tool was used to compare *Sc-dpy-10* and *Cel-dpy-10* and identified conserved Arg-to-Cys mutation sites (Madeira et al., 2022). The sequence of target genes in *S. hermaphroditum* were extracted from NCBI reference genome (BioProject: PRJNA982879; accession JAUCMV010000005). The gene structures were generated by Exon-Intron Graphic Maker 2012 v.4 (http://www.wormweb.org/exonintron). CRISPRscan was used to identify protospacer-adjacent motif (PAM) sites and design CRISPR RNA (crRNA) sequences (Table S1) (Moreno-Mateos et al., 2015). The Cas9 ribonucleoproteins were produced using protocols adapted from *C. elegans* CRISPR-Cas9 genome-editing (Wang et al., 2018; Wong et al., 2019). Briefly, to anneal guide RNAs, 2.5 µL of 100 µM of crRNAs and tracrRNA (the Alt-R system from IDT, Coralville, IA) are mixed at 1:1 ratio and incubated at 94°C for 2 minutes and cooled to room temperature. **For co-CRISPR in *S. hermaphroditum***, 1.5 µL of 100 µM of *Sh-unc-22* crRNA (co-CRISPR marker) and 1.5 µL of *Sh-daf-22* target gene crRNA (100 µM) are mixed with 3 µL of tracrRNA (100 µM). To prepare the injection solution, 3.4 µg/µL Ca9 protein and 45.9 µM of the annealed guide RNA are mixed. **In *S. carpocapsae* injection mixture**, 0.75 µM of single stranded (ss)DNA repair oligo (IDT) and 12% Lipofectamine®RNAiMax reagent (Invitrogen, Waltham, MA) was added to the injection mixture (Adams et al., 2019). The injection mixture was spun down at 15,000 rcf for 1 minute and incubated at room temperature for 10 minutes. The Cas9 RNP complex was then kept on ice and immediately delivered to *Steinernema* nematodes via gonadal microinjection.

### Gonadal microinjection of S. hermaphroditum and S. carpocapsae

***S. hermaphroditum*** young adult hermaphrodites (P0) were grown on *X. griffiniae,* rinsed in M9 buffer (per liter: 3 g KH_2_PO_4_, 6 g Na_2_ HPO _4_, 5 g NaCl, with 1 ml1 M MgSO_4_ added after autoclaving) to remove bacteria, and placed onto a 2% agarose pad mounted on #1, 50 x 22 mm glass coverslip (Corning, Corning, NY) immersed in Halocarbon oil 700 (Sigma, St Louis, MO). To create injection needles, borosilicate glass capillaries (World Precision Instrument, 1B120F-4: 4 in., OD-1.2 mm, ID-0.68 mm; filament, fire polish) were pulled using a pipette puller (P-87, Sutter Instruments, Novato, CA). A sharp opening was made on the tip of the needle. Cas9 RNP complex was loaded onto the needle and injected into the young adult gonad syncytia (Fig. 1B) using FemtoJet 4x (Eppendorf, Hamburg, Germany) at maximal (cleaning) pressure at approximately 87 psi. Injected nematodes were immediately washed in M9 buffer to remove injection oil, then recovered on a lawn of *Commamonas aquatica* (DA1877) on NGM agar plates at 25°C overnight, then isolated onto individual lawns of either *C. aquatica* or *X. griffiniae.* Twitching (*Sh-unc-22* phenotype) was screened in F1 progenies using 2% nicotine assay modified from (Gang et al., 2017; Moerman et al., 1988). Briefly, nematodes were screened in 2% nicotine (Sigma) and *Sh-unc-22* mutants were identified with a twitching phenotype (Video S1 and Video S2). Dumpy (*Sh-dpy-10* phenotype) were identified in F2 progenies.

**To microinject *S. carpocapsae* (All strain)**, IJs were surface-sterilized with 1% bleach, washed in sterile water, and incubated on *X. nematophila* (HGB800) lawns on lipid agar plates for 48 hours. Young adult females were injected using similar method as *S. hermaphroditum* and recovered on NGM agar with 2 µL of 1:10 diluted *E. coli* OP50 overnight culture. After incubating at 25°C for 3 hours, males were introduced to the agar plate for mating and incubated overnight at 25°C (Fig. S1). Gravid females were then isolated onto individual 0.5x lipid agar plates (per liter: 15 g bacto agar, 4 g nutrient broth, 2.5 g yeast extract, 2 gMcCl_2_ . 6H_2_O, 3.5 ml corn syrup, and 1 ml 5 mg/mL cholesterol in ethanol). On-target modification in F1 and F2 progeny were detected by genotyping and Sanger-sequencing.

### Genotyping and Sanger sequencing to confirm on-target genome editing

Individuals or a population of *Steinernema* nematodes were lysed in lysis buffer (50mM KCl, 10mM Tris pH8.0, 2.5mM MgCl_2_, 0.45% NP-40, 0.45% Tween 20, 0.01% gelatin; add 1 mg/mL Proteinase K immediately before use). To freeze-crack the nematodes and release genomic DNA, samples were incubated at -80°C for 15 minutes, 65°C for 10 minutes, then at 95°C for 15 minutes. Large insertions and deletions (indels) were detected by the presence and absence of PCR product using wild-type as a control on 1-2% agarose gel. Short indels in both *S. hermaphroditum* and *S. carpocapsae* were confirmed by Sanger sequencing of PCR product using genotyping primers designed by Primer 3 v.4.0 (Table S2). Homology-directed repair in *S. carpocapsae* was first detected by restriction digestion of the genotyping PCR product HindIII (New England Biolabs, Ipswich, MA), then confirmed by Sanger sequencing (Laragen, Culver City, CA).

### Co-CRISPR in S. hermaphroditum

Injected P0 animals were recovered as described above and isolated onto individual *X. griffiniae* lawns on NGM agar plates. F1 progenies with a twitching phenotype (with or without 2% nicotine treatment), and their hermaphroditic siblings were isolated to produce F2 self-progeny. To detect if F1 progeny has *Sh-daf-22* modification on target, eight F2 progeny were randomly chosen from F1 candidates (twitcher and twitcher’s siblings) to perform single-nematode *Sh-daf22* genotyping followed by Sanger sequencing. Once long and short indels were detected in F2 progenies by genotyping, their non-twitching (wild-type allele in *Sh-unc-22*) F2 siblings were chosen to homozygote the *Sh-daf-22* mutant alleles. Homozygous lines of *Sh-daf-22* mutants were maintained by cryopreservation (Cao et al., 2021).

### IJ (dauer larvae) entry assays

To perform IJ entry assay ***in vivo (in insecta)***, approximately 50-100 wild-type and *Sh-unc-22* IJs were used to infect each *Galleria* insect. Because *Sh-daf-22* showed a defect in establishing infection in *Galleria,* approximately 500 IJs (which may contain pre-IJs) were used to infect each *Galleria* insect at 25°C. IJs were white-trapped at room temperature (White, 1927). Nematode populations that emerged from the insect cadaver were resuspended in M9 buffer and transferred from filter paper onto NGM agar plates to be imaged to estimate percent IJ and population composition.

To perform IJ entry assay ***in vitro,*** approximately 200 axenic embryos were extracted and grown on Liver-Kidney agar seed with *X. griffiniae* lawn and incubated at 25°C (Murfin, Chaston, et al., 2012; Vivas & Goodrich-Blair, 2001). On days 8-11 of incubation, nematode populations at the edge of each petri-dish were resuspended in M9 buffer and the population composition was examined. Nematodes were imaged using SteREO Discovery.V20 (Zeiss, Jena, Germany). Images and videos were processed by ZenPro software and Fiji (Schindelin et al., 2012).

## Results

### Gonadal microinjection delivers CRIPSR-Cas9 ribonucleoprotein in *S. carpocapsae* and successfully modifies *Sc-dpy-10* homologue

*Steinernema carpocapsae* is the first *Steinernema* spp to publish a complete and annotated genome (Serra et al., 2019). I adapted the standard nematode microinjection technique based on that of *C. elegans* to deliver M9 buffer into the gonad of *S. carpocapsae*, in the pre-mated young adult female stage. In the initial trials, microinjection caused almost 100% death among animals, especially when injected nematodes were recovered on a lawn of symbiotic bacteria *X. nematophila* on lipid agar, a nutritional medium. This observation indicates that bacteria produce metabolites that kill the wounded nematodes. I further adapted the protocol to recover microinjected animals in M9 buffer and 1:10 diluted *E. coli* OP50 spots on a less nutritious NGM agar. This method improved animal survival rate to approximately 80% post-injection, enabling adult females to continue mating with wild-type males. In *C. elegans,* the collagen gene *dpy-10* causes cuticular morphological reorganization, resulting in a distinctive Dumpy or Dumpy/Roller phenotype (Levy et al., 1993). Specifically, a classical allele in *Cel*-*dpy-10,* Arg-to-Cys (R70C) substitution (*cn64*), causes a dominant Dpy/Roller phenotype (Levy et al., 1993). Similarly in *C. briggsae,* an R92C (*ot1210)* mutation in *Crb-dpy-10* is a dominant roller and is phenotypically consistent to *Cel-dpy10* cn64 (Toker & Hobert, 2022)*. S. carpocapsae* has one homologue of *dpy-10* (L596_943991 or SC.X.g3037, Fig. 2A) which showed 62.1% sequence identity with *Cel-dpy-10.* The *Sc-dpy-10* locus is linked to X chromosome which makes it an ideal target for genome-editing in a XX/XO dioecious species, because it facilitates genotyping and phenotyping in haploid F1 hemizygous male progeny. To create a mutation in *S. carpocapsae* that is similar to *Cel-dpy-10* R70C (*cn64*) and *Cbr-dpy-10* R92C (*ot1210*), I used a Multiple Sequence Alignment tool to find *Sc-dpy-10* Arg75 as a conserved residue (Fig. 2B). I targeted two PAM sites (both positive and negative strand) flanking *Sc-dpy-10* Arg75 (Fig. 2C). A short DNA repair template is designed to introduce a C-to-T transition that causes an R75C missense mutation. In addition, eight silent mutations are introduced to create a HindIII restriction digestion site to facilitate genotyping (Fig. 2C).

**Figure 2:**
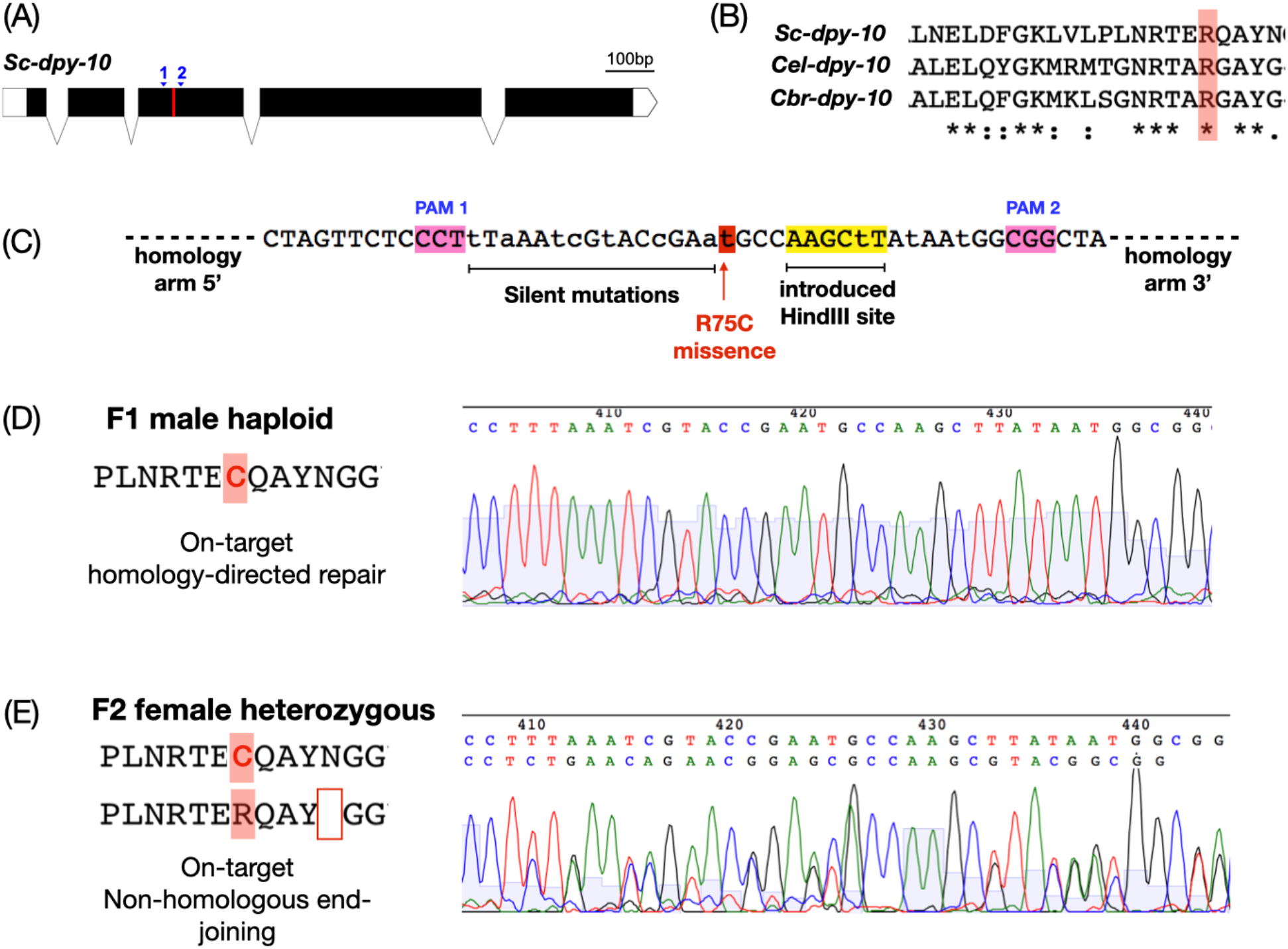
CRISPR-Cas9 based technique creates on-target and heritable modifications of *Sc-dpy-10.* (A): *Sc-dpy-10* gene structure. Blue triangles denote PAM sites 1 (non-coding) and 2 (coding). Red stripe denotes Arg75. (B): Multiple sequence alignment of *S. carpocapsae, C. elegans,* and *C. brigssae* Dpy-10 homologues showed conserved Arg75 residue in Sc-Dpy-10. Asterisk (*) denotes identical amino acids; colon (:) denotes conservative mutations; dot (.) denotes semi-conservative mutations; space ( ) denotes non-conservative mutations. (C): ssDNA repair template is designed to introduce mutations (lower case letters) including R75C missense (red) and eight silent mutations. A HindIII restriction digestion site is introduced to facilitate genotyping. (D): Sanger sequencing chromatogram shows F1 male (X-linked R75C) progeny with on-target modification by homology directed repair. (E): Sanger sequencing chromatogram shows F2 heterozygous female animal with one R75C allele by HDR and one N79 deletion allele by non-homologous end joining (NHEJ), confirming both F1 parents carry heritable *Sc-dpy-10* mutation alleles from on-target modifications.

Out of the thirty-six P0 adult females I injected, thirty animals survived, and six P0 lines produced viable F1 animals with no obvious Dumpy or Roller phenotypes. Alternatively, I used single F1 animal genotyping to identify on-target modification of *Sc-dpy-10* sequence. One of the six lines (designated line 18) showed HindIII digestion on the 792 bp of PCR-amplified product and is further confirmed by Sanger sequencing as a successful introduction of R75C along with eight silent mutations (Fig. 2D). The R75C point mutation and an N79 deletion at PAM 2 were both identified among F2 progeny of Line 18 (Fig. 2D, Fig. S1B), confirming both homology-directed repair (HDR) and non-homologous end joining (NHEJ) occurred, and these mutations are heritable for two generations. Various on-target modifications were also detected by genotyping F2 progeny (Fig. S1). My initial attempts of microinjecting CRISPR-Cas9 in *S. carpocapsae* created on-target and heritable gene modifications, however, I did not observe the expected phenotype of Dumpy or Roller in these mutant animals, suggesting this allele may be specific to *Caenorhabditis* nematodes.

### On-target modification of *Sh-dpy-10* homologue causes Dumpy phenotypes in *S. hermaphroditum*

I adapted similar Cas9 RNP microinjection techniques in *S. hermaphroditum*. I first discovered that *S. hermaphroditum* post-injection survival increased from approximately 70% (on *X. griffiniae* on NGM agar plates) to 95% (on *C. aquaticus*), therefore adapted the protocol to recovering injected animals on the *C. aquaticus* lawn. The *Sh-dpy-10* sequence was predicted by BlastP using Cel-Dpy10 as a reference. Gene structure showed five exons that translates into a 344 amino acid peptide (Fig. 3A). Facilitated by self-fertilization, two homozygous F2 recessive mutant lines were identified and isolated by visible Dumpy phenotypes (MCN0001 and MCN0002, Fig. 3E, 3F, and 3G). Genotyping and Sanger sequencing confirmed that *mc0001* carries a *Sh-dpy-10* homozygous allele with modifications at both PAM 1 (Exon 1) and PAM 2 (Exon 2), introducing an early stop codon after 11 amino acids (null mutant, Fig 3B and 3D). The allele *mc0002* showed a modification at PAM 2 with 39 base pairs deletion in Exon 2 coding region, causing a truncated Dpy-10 protein (Fig. 3C and 3D). However, neither line could recover from cryopreservation, suggesting they may have a defect in cryobiology. Overall, both *Sh-dpy-10* null and in-frame deletion mutant lines showed heritable and visible phenotype as expected.

**Figure 3:**
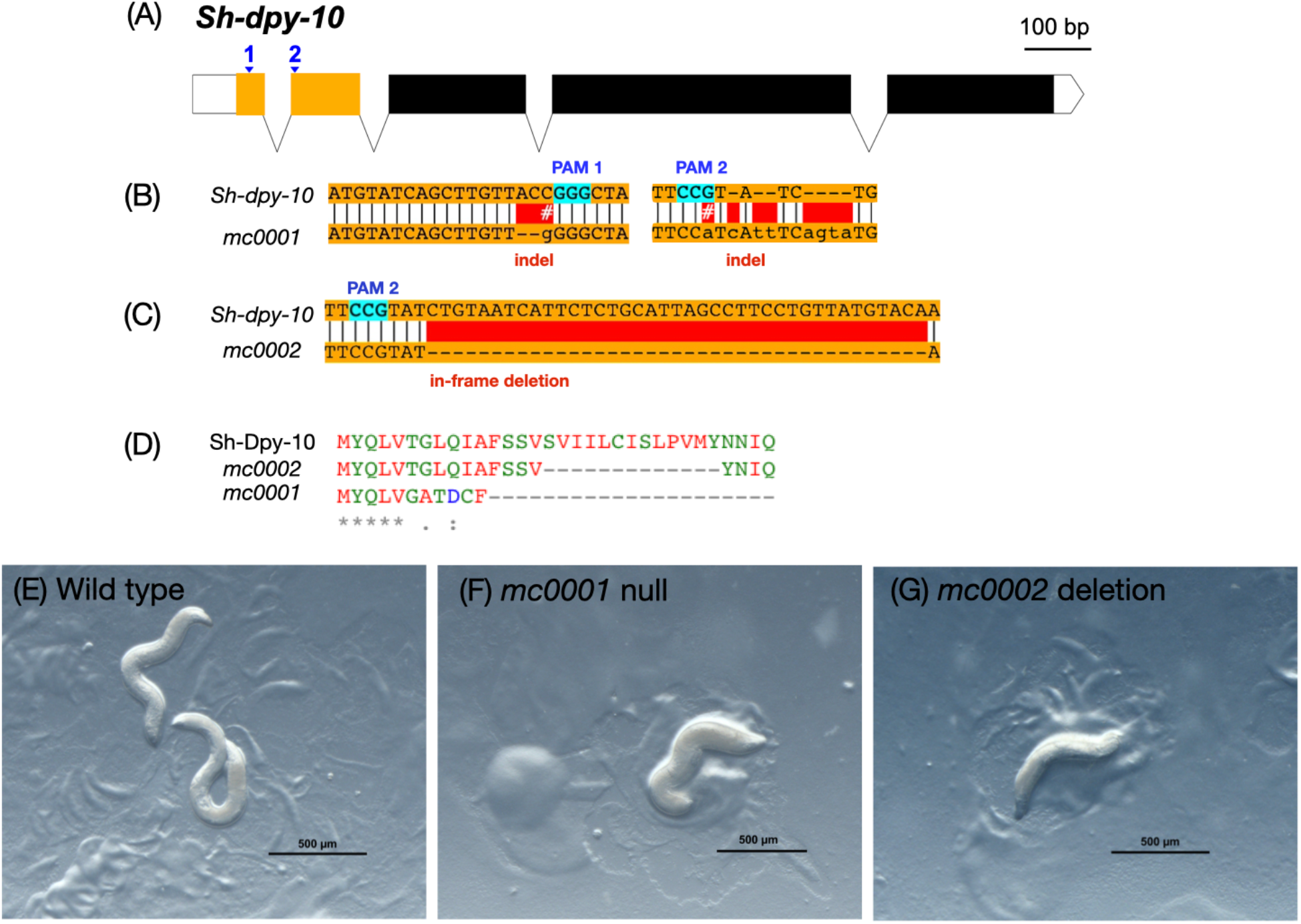
Cas9 nucleoprotein delivered by gonadal microinjection created *S. hermaphroditum dpy-10* mutants with expected phenotypes. (A): *Sh-dpy-10* homologue gene structure. Blue triangles denote PAM sites 1 and 2. (B and C): Sanger sequencing shows on-target modifications of *Sh-dpy-10,* creating alleles *mc0001* (B) and *mc0002* (C). (D-F): Young adults of wild-type *S. hermaphroditum* (E), *mc0001* with an early stop codon in *Sh-dpy-10* (F), and *mc0002* with13 aa deletion in *Sh-dpy-10* (G). Scale bar = 500 µm.

### *Sh-unc-22* mutation creates conditional dominant twitching phenotypes

To seek for a dominant allele that produces a consistent and visible phenotype as a co-CRISPR marker, I targeted *Sh-unc-22,* predicted to encode a muscle gene termed Twitchin protein in free-living and parasitic nematodes (Gang et al., 2017; Hellekes et al., 2023; Moerman et al., 1988). In nematodes, UNC-22 is crucial in maintaining muscle contraction cycles. Mutants of *unc-22* shows spasmodic muscle twitching. Such a twitching phenotype is dominant in response to cholinergic agonists, such as nicotine and the anti-parasite paralyzing drug levamisole (Gang et al., 2017; Lewis et al., 1980). Sh-Unc22 was located as a hypothetical protein QR680_003339 (Accession: KAK0400070.1), which shares 95% coverage and 68.9% amino acid sequence identity with Cel-Unc22. The gene encoding *Sh-unc-22* is predicted to contain 39 exons, in which Exon 33 is highly conserved (67.23% amino acid sequence identity with Cel-Unc22); therefore it was chosen as the targeted region for editing (Fig. 4A). Approximately 5-10% of the injected P0 animals produced F1 progeny with twitching in 2% nicotine as an expected phenotype (Fig. 4D, Video S1 and S2). Genotyping of F1 and F2 twitching progeny confirmed on-target modification in all five PAM sites, creating diverse mutant alleles (Fig. 4E, Fig. S2). I maintained two homozygous and healthy mutant lines with strongly twitching phenotypes. Each one showed in-frame deletion, insertion, and substitution of amino acids at PAM 3 and PAM 5 (Fig. 4B and 4C). Mating tests confirmed *Sh-unc-22* (*mc0003)* is an X-linked conditionally dominant allele (Fig. S3): the heterozygous hermaphrodites (X*^unc-22^*/X^+^) twitch exclusively in 2% nicotine, but not in M9 buffer, while homozygous hermaphrodites (X*^unc-22^*/X*^unc-22^*) twitches under both conditions. Homozygous lines of *Sh-unc-22* animals are healthy and could be recovered after cryopreservation.

**Figure 4:**
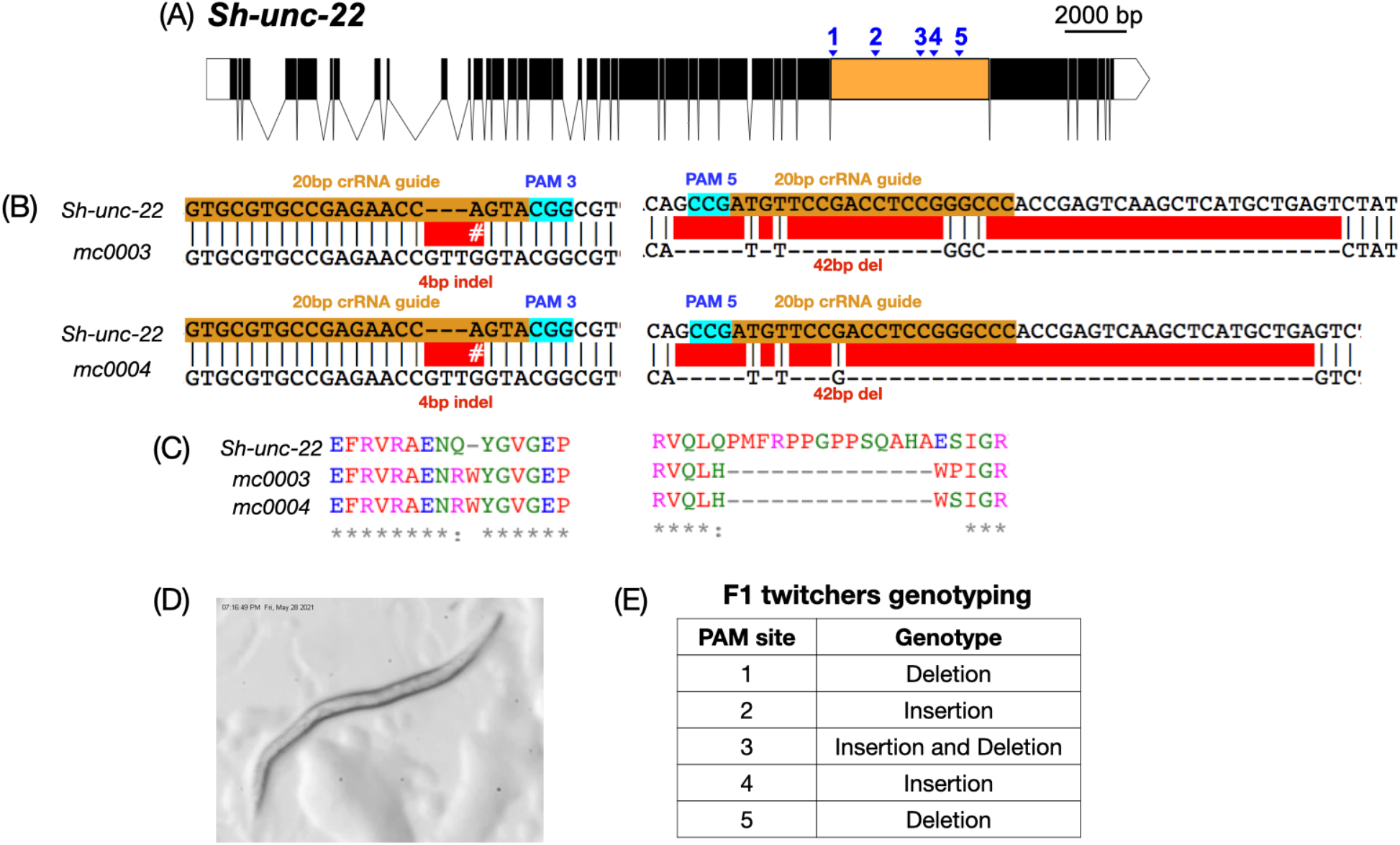
On-target modifications of *Sh-unc-22* homologue create conditional dominant alleles causing a twitching phenotype. (A): Gene structure of *Sh-unc-22* and targeted PAM sites 1-5. Blue triangles: PAM sites; orange block: highly conserved Exon 33. (B): F2 twitchers carry homozygous alleles of *unc-22* mutations with on-target modifications at PAM 3 and PAM 5. (C): Twitchers showed in-frame modifications of amino acids, including insertions, deletions, and substitutions. (D): A representative juvenile nematode with *unc-22* mutation. (E): A table summarizes the detected modifications at each PAM site.

### *Sh-unc-22* as a Co-CRISPR marker to target *Sh-daf-22*

The modification of *Sh-dpy-10* and *Sh-unc-22* both showed approximately 5-10% of injected P0 animals with editing in their germlines, out of which approximately 3-30% of the F1 animals have heritable mutations shown by phenotype. With this efficiency, to create a homozygous recessive allele without an immediately visible phenotype, it would take genotyping and sequencing approximately 1,280-4,800 F2 animals to produce 1-2 mutant lines. I propose to further use conditionally dominant allele *Sh-unc-22* as a co-CRISPR marker in *Steinernema*. As a proof of concept, I chose a second target locus encoding a highly conserved homologue, Sh-Daf-22, a core enzyme required for nematode pheromone ascaroside biosynthesis (Ludewig & Schroeder, 2013). *S. hermaphroditum* has one homologue of Daf-22 (hypothetical protein QR680_000661; Accession KaK0394264.1), containing 536 amino acids that has 96% coverage and 78% identity with Cel-Daf-22. Transcriptomic analysis showed seven exons (Fig. 5B). I co-targeted *Sh-unc-22* PAM 3, PAM 5 (Fig. 4), and *Sh-daf-22* PAM 1 (Exon 2) and PAM 2 (Exon 3) (Fig. 5A and 5B). As expected, approximately 5-10% of injected P0 animals produced twitching F1 progeny, indicating Cas9 RNP had access to edit germ cells in these P0 animals. Within the lines of *Sh-unc-22* modification, I examined non-twitching F2 progeny derived from twitching F1 animals and approximately 25% of the animals carried on-target mutations at *Sh-daf-22* PAM sites. I also examined the F2 progeny from siblings of F1 twitchers (P0 germline with Cas9 RNP access). The on-target mutations of *Sh-daf-22* were approximately 20%, suggesting that the editing of *Sh-unc-22* and *Sh-daf-22* were independent in these lines. This feature makes *Sh-unc-22* (X-linked) a useful injection marker even if the second target is on the same chromosome. I genotyped sixteen F1 lines from two P0 animals (128 animals from F2 generation) to secure three mutant lines of heritable and homozygous alleles in *Sh-daf-22* (Fig. 5B): a 21 bp insertion causing in-frame amino acid modifications at PAM site 1 (*mc0005*); a 171 base pair deletion at PAM site 1 (spanning Exon 2 to Exon 3) causing a frame-shift after 42 amino acids and an early stop codon after 75 amino acids (*mc0006*); and a truncated protein (total 251 amino acids) with deletions at both PAM 1 and PAM 2 spanning from Intron 1 to Intron 5 *(mc0007)*. The successful enrichment of *Sh-daf-22* mutant alleles demonstrates *Sh-unc-22* as a useful co-CRISPR marker in *S. hermaphroditum*.

**Figure 5:**
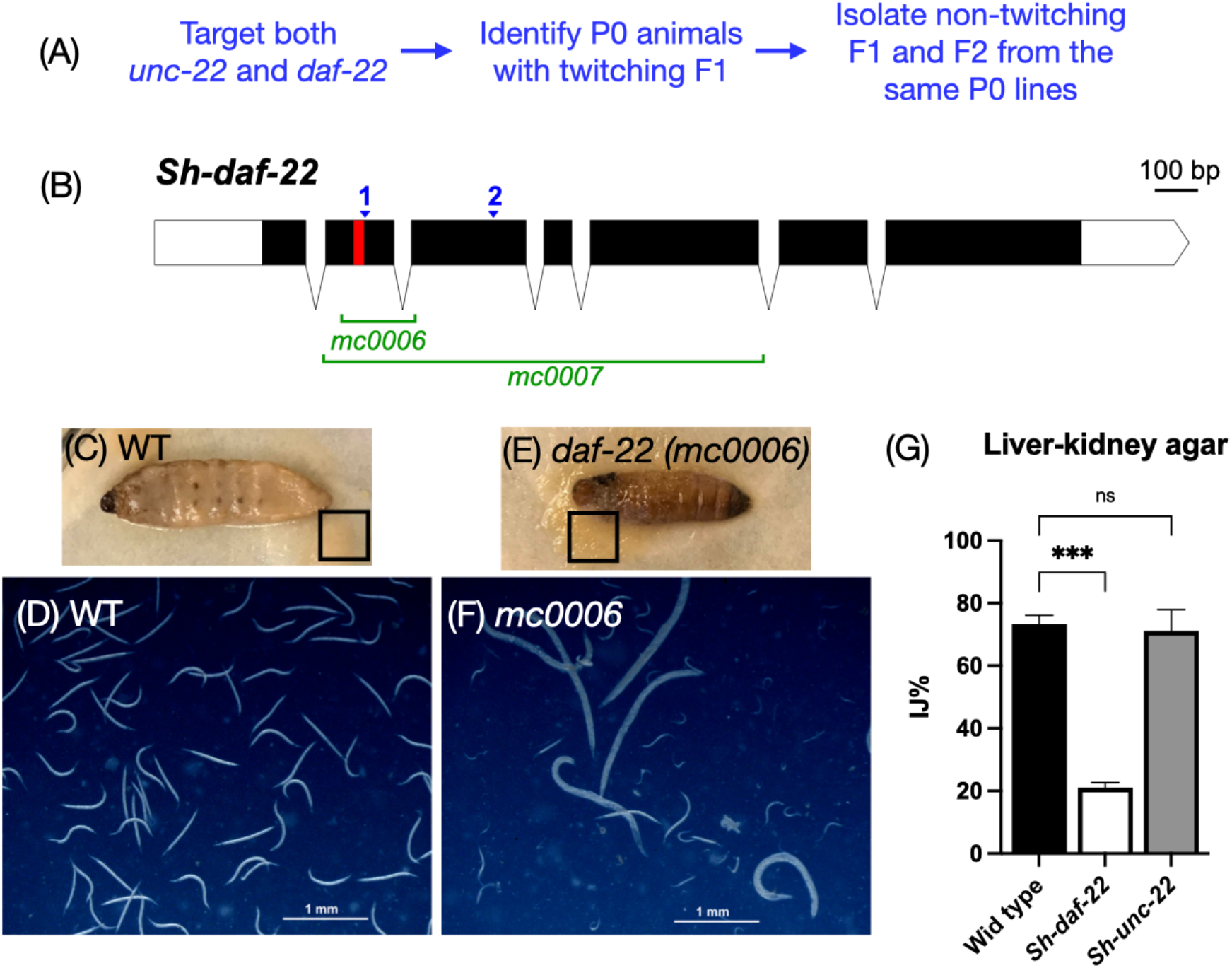
*Sh-unc-22* as a co-CRISPR injection marker to facilitate the modification of a second gene, *Sh-daf-22*. (A): An experimental flowchart of genome-editing by co-CRISPR marker. (B): Gene structure of a homologue of *Sh-daf-22* as a second target. Blue triangles denote PAM sites 1 and 2; the red band denotes 21 base pairs insertion in allele *mc0005*. Green brackets denote deletions in *mc0006* and *mc0007*. (C-H): Mutant allele of *Sh-daf-22* showed dauer (IJ) entry defect *in vivo*. Nematodes emerged from *Galleria* insect cadaver (indicated by the black box) infected with wild-type (C), *Sh-daf-22* mutant (E) were resuspended in M9 buffer and examined by microscopy. Emerged wild-type (D) are nearly 100% IJs. Emerged *Sh-daf-22* (F) nematodes are primarily adults and juveniles, with approximately 10% IJs. (G): Quantification of percentage of IJs in the nematode population on nutrient-rich liver kidney agar.

The *daf-22* gene encodes a homologue of human sterol carrier protein SCPx, which catalyzes the final step in peroxisomal fatty acid β-oxidation (Butcher et al., 2009). In nematodes, *daf-22* processes the small molecule ascarosides (nematode pheromone) by cleaving and shortening the fatty acid side chains (Butcher et al., 2009). A signature phenotype of the *daf-22* mutation in nematodes is a defect in dauer larvae (paralleled stage of IJ) development, a process dependent on dauer pheromone production (Butcher et al., 2009; Cohen et al., 2022; Golden & Riddle, 1982; Markov et al., 2016). Entomopathogenic nematodes also produce ascarosides that affect IJ development and dispersal behavior (Choe et al., 2012; Kaplan et al., 2012; Noguez et al., 2012; Roder et al., 2019), however, the composition and function of ascarosides in EPN is limited, and the function of *daf-22* is unknown, due to the lack of genetic tools. As a proof of concept, I performed *in vivo* dauer (IJ) entry assay by infecting *Galleria* insect larvae (waxworm) with *S. hermaphroditum* IJs of wild type and *Sh-daf-22 (mc0006)*. In a normal life cycle, entomopathogenic nematodes reproduce in the infected insects and emerge as IJs in response to environmental cues including pheromones (Roder et al., 2019). I collected and examined the population of nematodes that emerged from the *Galleria* insects (Fig. 5C and 5E). As expected, the wild-type *S. hermaphroditum* showed 100% IJs in the emerged population (Fig. 5D). In contrast, *Sh-daf-22* null mutant (*mc0006*) showed an IJ developmental defect, with 10%-50% IJs and most of the population being adults and juveniles in the emerging population (Fig. 5F). The IJ developmental defect is independent of the *Sh-unc-22* co-CRISPR marker (Fig. S4 and S4B). This IJ developmental defect is quantifiable on liver-kidney media (made of ground beef liver and kidney), mimicking a nutrient-rich insect cadaver (Fig. 5G and Fig. S4). These data show that *daf-22* mutant has an IJ developmental defect in the infected insect, a relevant host environment in the nematode natural life cycle. This exemplifies *S. hermaphroditum* as a promising model to study gene functions in the relevant ecological context.

## Discussion

This research established a CRISPR-Cas9 based protocol in entomopathogenic nematodes and showed successful gene editing in *S. carpocapsae* and *S. hermaphroditum.* In *S. hermaphroditum,* modification of *Sh-dpy-10* and *Sh-unc-22* both produced progeny with expected and visible phenotypes, dumpy and twitching, respectively. Using *Sh-unc-22* as a co-CRISPR injection marker, mutations were enriched in a second gene, *Sh-daf-22,* which showed a IJ developmental defect within infected insects. In *S. carpocapsae,* homology directed repair (HDR) has been observed, suggesting such mechanism is present in *Steinernema.* Overall, this work is a proof of concept that *Steinernema* nematodes are highly tractable. As this manuscript is being prepared, others adopted this protocol and claimed it was successful in an independent work, proving this method to be consistent and generalizable (Schwartz et al., 2023). The molecular tool development in entomopathogenic nematodes, exemplified by CRISPR-Cas9 genome editing, opens new opportunities to investigate gene functions in an animal’s natural habitat and complements our knowledge that may not be feasible to acquire from other genetic models.

### *Steinernema hermaphroditum* has a syncytial gonad that facilitates transformation

In *S. hermaphroditum,* the microinjection is not reliant on the use of transfection agent lipofectamine, proving that *S. hermaphroditum* has a syncytial gonad in which nuclei of germ cells are accessible to Cas9 RNP editing. One preliminary trial using lipofectamine in *S. hermaphroditum* gonadal injection reduced post-injection survival rate from >90% to <50%, suggesting lipofectamine may be toxic to the animals. In *S. carpocapsae,* I used lipofectamine in the successful cases, but it’s unclear whether transfection is absolutely required, because the success rate of editing was low in this species regardless. The microinjection equipment for *Steinernema* is the same as that of *C. elegans,* and the techniques and workflow is very similar. Therefore, it is highly promising to adapt other established techniques in *C. elegans* transformation in *Steinernema* spp.

### Symbiotic bacteria harm the wounded nematodes post injection

The survival rate of injected animals is highly dependent on the bacterial species on which the nematodes are recovered. In general, the survival rate of wounded animals is significantly lower on their native symbiont in comparison to other bacteria, such as *E. coli* and *C. acquaticus.* Injected *S. carpocapsae* almost had 100% death if they were recovered on their symbiont*X. nematophila,* while recovering on *E. coli* OP50 significantly increased their survival. Wounded *S. hermaphroditum* survival rate also increased when nematodes were recovered on *C. acquaticus* in comparison to those on symbiont *X. griffiniae.* These observations suggest that *Xenorhabdus* bacteria could be toxic to their host nematodes under specific conditions, possibly when the nematodes have a damaged cuticle, or when they are under highly stressful situations such as high-pressure microinjection. *Xenorhabdus* bacteria produce secondary metabolites that function as toxins and antimicrobials to protect the infected insects from other microbes, a phenomenon termed as defensive symbiosis (Murfin et al., 2019). It is unclear how *Steinernema* immune system tolerates their symbiont, and if such pathway is the same as nematode immunity in response to a pathogen. Pursuing these questions will be require further genetic tool development in EPN.

### Homology-based repair in *Steinernema* CRISPR

I directly injected *in vitro-*assembled Cas9 RNP complexes in the gonad of P0 nematodes, using linear ssDNA with short homology arms (35-50 bp) as repair templates. In *C. elegans,* this technique was proved to yield homology directed repair (HDR) at a high frequency (Paix et al., 2015, 2017). In *S. carpocapsae,* I detected successful HDR at a very low frequency (one F1 line out of from 300 injections of P0 animals, <0.3%). In *S. hermaphroditum*, injection of more than 100 P0 animals using this technique did not yield a detectable allele of HDR. Other techniques may also be required to improve HDR efficiency in these species.

### Phenotypes of *Steinernema* mutants

As a proof of concept, I strategized to modify conserved genes that could serve as genetic markers in *Steinernema*. I first targeted *dpy-10* homologues, a cuticular collagen gene, known to control body shape and size in *C. elegans,* such as Dumpy and Roller (Levy et al., 1993; Toker & Hobert, 2022). *Sc-dpy-10* R75C mutation did not cause a dominant dumpy/roll phenotype as it is in *Caenorhabditis* species. Null and deletion mutations in *Sh-dpy-10* showed a mild Dumpy phenotype in approximately 10-20% of J4 and adult stages. These observations suggest that the function of *dpy-10* in cuticular re-organization may not be identical among *Steinernema* and *Caenorhabditis spp.* In *C. elegans*, *dpy-10* was shown to regulate *C. elegans* body stiffness, resistance to osmotic stress, and innate immune response (Fechner et al., 2018; Wheeler & Thomas, 2006; Zhu et al., 2023). Consistent with *Cel-dpy-10* functions, I found neither *Sh-dpy-10* mutant lines can be recovered from cryopreservation, a defect in nematode cryobiology that is relevant to their resistance to high osmolarity (Gade et al., 2020). The conserved and unique roles of *dpy-10* in *Steinernema,* specifically in response to its insect host and symbiotic bacteria, will be explored as these nematodes as they are established into new genetic models.

The in-frame deletions of *Sh-unc-22* caused twitching phenotypes in *S. hermaphroditum*, suggesting *unc-22* function is highly conserved among diverse nematode species. Similar to *Cel-unc-22,* heterozygous *Sh-unc-22* animals twitch in 2% nicotine (conditionally dominant), while homozygous animals twitch with and without nicotine (Moerman et al., 1988; Moerman & Baillie, 1979), making this allele a useful co-CRISPR marker that has proven to be successful in *S. hermaphroditum.* The *Sh-unc-22* mutants could be propagated through insects (Video S1 and S2; Fig. S4) and they recover from cryopreservation. For these reasons, this co-CRISPR marker may be further adapted to other nematode species. UNC-22 is associated with calcium release and calcium-sensitive contraction in nematodes, and *unc-22* mutants are resistant to levamisole, a paralyzing reagent used for treating roundworm infections in humans and animals. Despite its importance in parasitology, current knowledge in levamisole-resistance is highly dependent on the studies in non-genetically tractable parasitic nematodes and free-living nematode *C. elegans* (Sangster et al., 2005). This work shows that EPN, with parasitic life cycle, genetic tractability, and no ethical concerns, may be used as a new model to explore parasite-specific drug-resistant genes.

Our current knowledge in the composition, identity, and functions of ascarosides in EPN is very limited, because of the lack of genetic tools in these nematodes. This work shows that *Sh-daf-22* null mutant is defective in IJ (dauer) entry, which is consistent with phenotype of *Caenorhabditis* and *Pristionchus daf-22* mutants. To explore the function of *Sh-daf-22* in *S. hermaphroditum* IJ development, I first adapted a protocol from *C. elegans* dauer entry assay: I extracted pheromone from nematode liquid culture, which was then added exogenously onto nutrient-limiting (NGM-based) agar to induce IJ formation (Golden & Riddle, 1982). This protocol showed pheromone has a minor effect in IJ entry (data not shown) in comparison to food and temperature, either because liquid culture does not contain sufficient amounts of pheromone, or because the *in-vitro* conditions in this experiment are irrelevant to the development of this nematode species. In contrast, the *in-vivo* assays using insects or *in-vitro* assay using nutrient-rich liver-kidney agar showed drastic differences of IJ developmental defect from *Sh-daf-22* mutant, suggesting this gene may function differently *in vivo* in comparison to the standard laboratory conditions established for *C. elegans* experiments. Extensive study is required to fully understand the function of these genes in the natural life cycle of *Steinernema*.

## Acknowledgements

Paul W. Sternberg, Heidi Goodrich-Blair, Chieh-Hsiang Tan, and Jennifer Heppert provided helpful discussions for this project. Lorrayne Serra, Ali Mortazavi, Adler R. Dillman, Hillel T. Schwartz, and Erich M. Schwarz shared their knowledge on *S. carpocapsae* and *S. hermaphroditum* genomes and blasting target genes. Sally Adams and Pire DaSilva shared their knowledge in liposome-based delivery method. Wan-Rong Wong and Heenam Park shared their expertise in *C. elegans* CRISPR-Cas9 techniques. Carly Myers assisted in performing *in-vivo* IJ entry assays. Elin Larsson, Carly Myers, and Phoebe Lostroh helped proof-read the manuscript. Paul W. Sternberg and Richard M. Murray provided funding and space. Margaret J. McFall-Ngai shared equipment.

## Funding

This research is supported by National Science Foundation (NSF) Enabling Discovery through Genomics (EDGE) grant 2128267 (to P.W. Sternberg); National Institutes of Health (NIH) Ruth L. Kirschstein National Research Service Award (NRSA) Individual Postdoctoral Fellowship F32 5F32GM131570 (M.C.), the Resnick Sustainability Institute (California Institute of Technology), and Carnegie Institution for Science endowment.

## Notes

### Competing Interest Statement

The authors have declared no competing interest.

