## Supplemental Data for "CRISPR-Cas9 genome editing in *Steinernema* entomopathogenic nematodes"

**Video S1:** Homozygous *Sh-unc-22* mutant allele (*mc0003/mc0003*) causes IJ twitching in water-trap.

**Video S2**: Wild-type IJs emerged from a *Galleria* insect and kept in a water-trap.

**Table S1**: A list of guide RNAs used in this research.

| Target gene | crRNA sequence | PAM site |
|---|---|---|
| Sc-dpy10-cr1- | TGGCGCTCCGTTCTGTTCAG | Pam 1 |
| Sc-dpy10-cr2+ | GAGCGCCAAGCGTACAACGG | Pam 2 |
| Sh-dpy10-cr1+ | AGATGTATCAGCTTGTTACC | Pam 1 |
| Sh-dpy10-cr2- | GCAGAGAATGATTACAGATA | Pam 2 |
| Sh-unc22-cr1- | GCGCCTCCATTGTCAGTGGG | Pam 1 |
| Sh-unc22-cr2- | AGGAGCGGGACGGTCGATGA | Pam 2 |
| Sh-unc22-cr3+ | GTGCGTGCCGAGAACCAGTA | Pam 3 |
| Sh-unc22-cr4- | TGGCATGGGGCTGGCAGCAT | Pam 4 |
| Sh-unc22-cr5- | GGGCCCGGAGGTCGGAACAT | Pam 5 |
| Sh-daf22-cr1+ | GCGACTGTTGGATACCTCTT | Pam 1 |
| Sh-daf22-cr2- | TGGAGAAGGGAAGAGACCAT | Pam 2 |

**Table S2**: A list primers used in this research.

| Gene | Short name | Sequences |
|---|---|---|
| Sc-dpy10_F | oMC86f | CGCGCTTCTTACCAGTTGAC |
| Sc-dpy10_R | oMC87r | GGGAGAGGTTCCTGGAGTTC |
| Sh-dpy10_F | oMC102f | CTCGAAGATCAAAGGCAAGC |
| Sh-dpy10_R | oMC103r | GCGGAAGGTCTATGCTTACG |
| Sh-unc22-Pam1_F | oMC119f | GCGAAGGTGAACAAGCTGAT |
| Sh-unc22-Pam1_R | oMC120r | CATCTGTGCGTCTGATGGAT |
| Sh-unc22-Pam2_F | oMC121f | CTATGTTTGAAGCCCCGAAA |
| Sh-unc22-Pam2_R | oMC122r | CTCCAAATTTCGAACCGTGT |
| Sh-unc22-Pam3_F | oMC123f | GGCAACCAGGAGATTTACGA |
| Sh-unc22-Pam3_R | oMC124r | GGTTTCCAGAGGTTCACCAA |
| Sh-unc22-Pam4_F | oMC125f | ATATGGCAATGCTCACACGA |
| Sh-unc22-Pam4_R | oMC126r | TCAAGCACTTCTCCACGATG |
| Sh-unc22-Pam5_F | oMC127f | CCGAATCCTCGGCTACAATA |
| Sh-unc22-Pam5_R | oMC128r | ATCTTGATGGGCGATGTAGG |
| Sh-daf22_F | oMC129f | CAAGTTTGTCAAGCCCCAAT |
| Sh-daf22_R | oMC130r | GGCTTGGTTCCTAAACGACA |

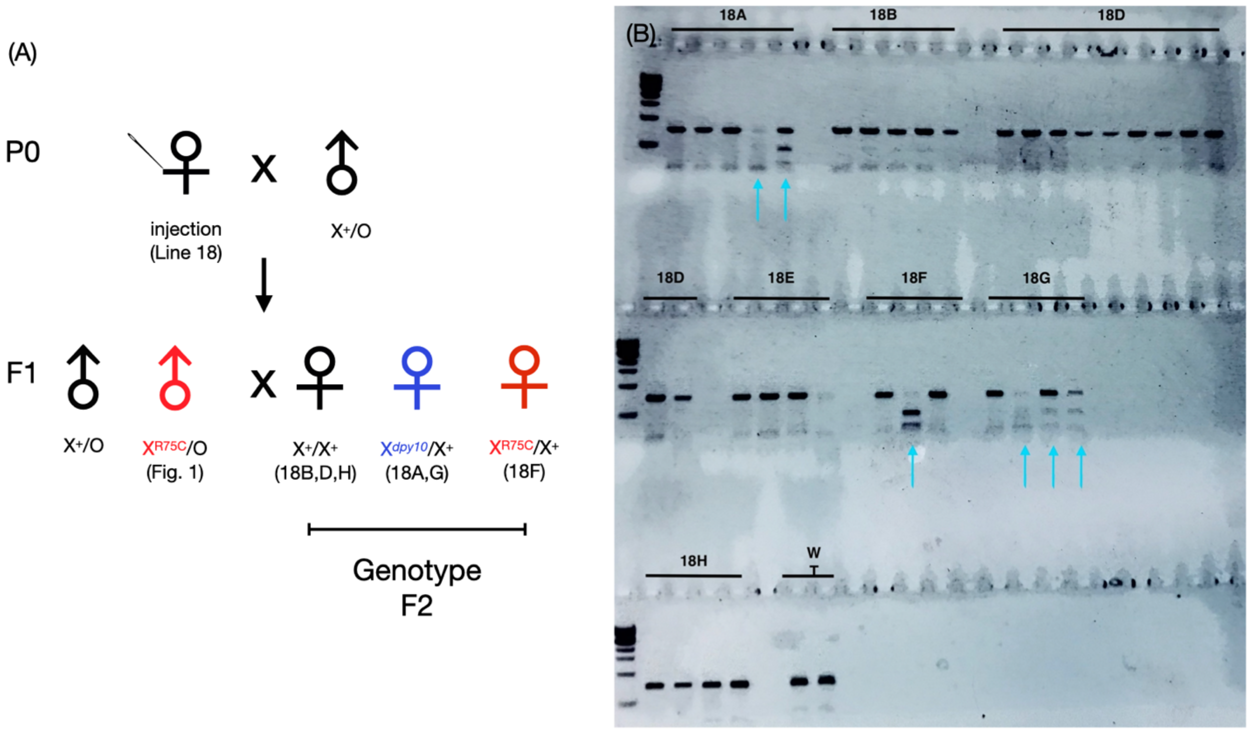


**Figure S1**: Single nematode genotyping of F2 progeny showed heritable on-target modification, producing *Sc-dpy-10* R75C and other alleles. (A): Schematic diagram of Line 18 with successful *Sc-dpy-10* modifications. The injected P0 female was mated with wild-type males. R75C mutation (red) was confirmed in F1 male (see Fig. 1). F2 progeny was produced by siblings mating. The genotypes of F2 (see Fig. S1B) confirmed multiple alleles created by CRISPR-Cas9 editing, including R75C (red) and others by NHEJ (blue). (B): Single F2 animal genotyping from Line 18 using PCR-amplification (792 bp band). HindIII digestion of *Sc-dpy-10* R75C allele produces two bands. Line 18A, 18B, 18D, 18E, 18F, 18G, 18H are seven F1 females that were mated with F1 male siblings. Each lane indicates genotyping and HindIII digestion from one F2 animal produced by designated F1 females. Blue arrows indicate a modification of *Sc-dpy-10:* a missing band indicates a deletion, two bands indicate HindIII digestion. WT: wildtype genotyping and HindIII digestion.

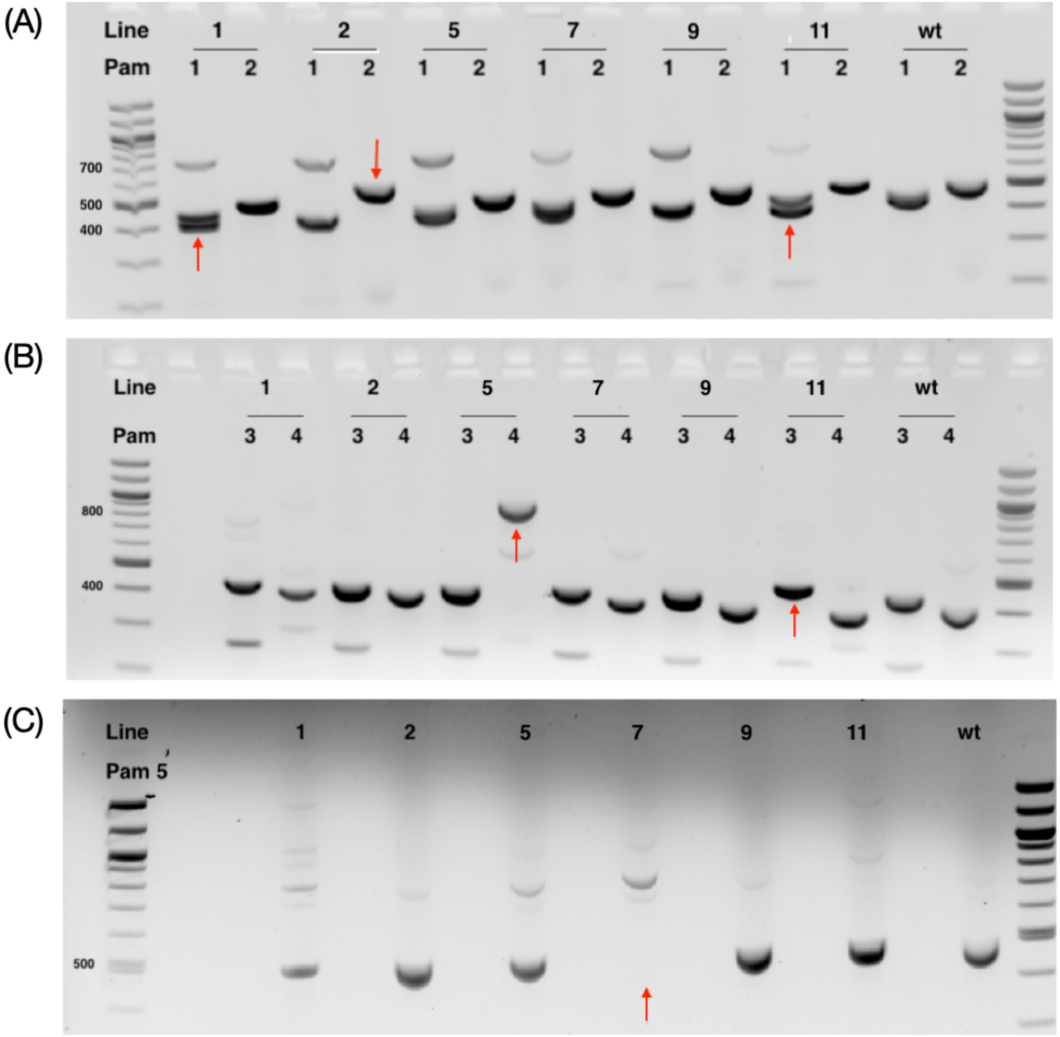


**Figure S2**: CRISPR-Cas9 targeting *Sh-unc-22* produces diverse modifications at five PAM sties. In one round of microinjection of twenty P0 adult hermaphrodites, eleven F1 lines were isolated from the same P0 line based on their twitching phenotype. Six of them produced live F2 lines (Line 1, 2, 5, 7, 9, and 11, shown on the gel). Representatives of F2 progenies of each of these six lines are genotyped by PCR amplification of 400-500 bp (wild-type band) around PAM sites 1 and 2 (A); PAM sites 3 and 4 (B); PAM site 5 (C). Overall, twitchers from five out of six lines showed at least one locus of on-target modification (indels) by genotyping.

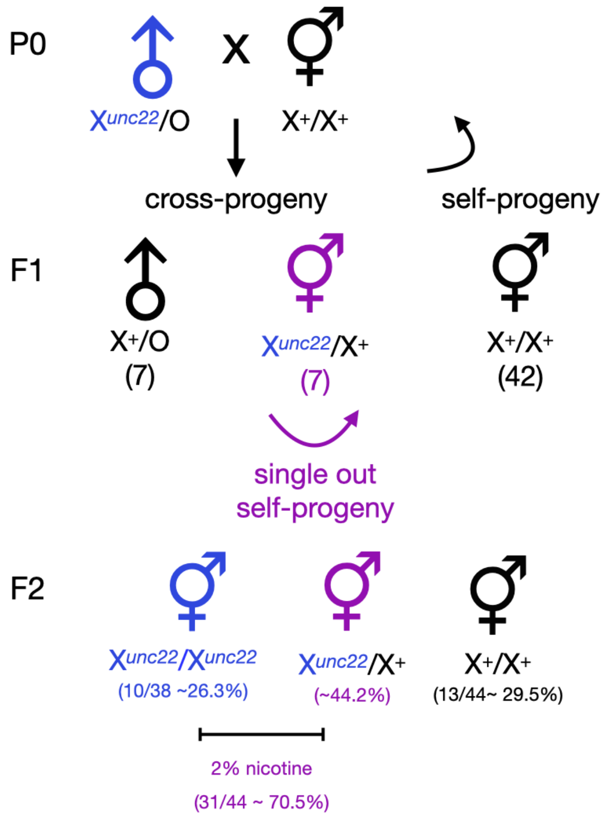


**Figure S3**: *Sh-unc-22* in-frame modification causes a dominant twitching phenotype in 2% nicotine. A mating test is used to confirm *Sh-unc-22* allele (*mc0003*) is an X-linked and conditionally dominant allele. P0 males from the Sh-unc-22 mutant line were mated with wild-type hermaphrodite. Among F1 progeny, 42 hermaphrodites do not twitch in 2% nicotine (self-progeny); 7 hermaphrodites twitched in 2% nicotine (conditionally dominant and cross-progeny), and 7 males do not twitch (*Sh-unc-22* is X-linked). Heterozygous F1 hermaphrodites were singled onto individual petri-dishes. F2 self-progeny showed Mendelian ratio of segregation. Blue: homozygous mutant that twitches with or without nicotine. Purple: heterozygous animals twitching in 2% nicotine only. Black: homozygous wild-type animals that do not twitch with or without nicotine. A cross 'x' denotes mating. Number of F1 progeny and percent population of F2 progeny in each category is indicated in the parenthesis.

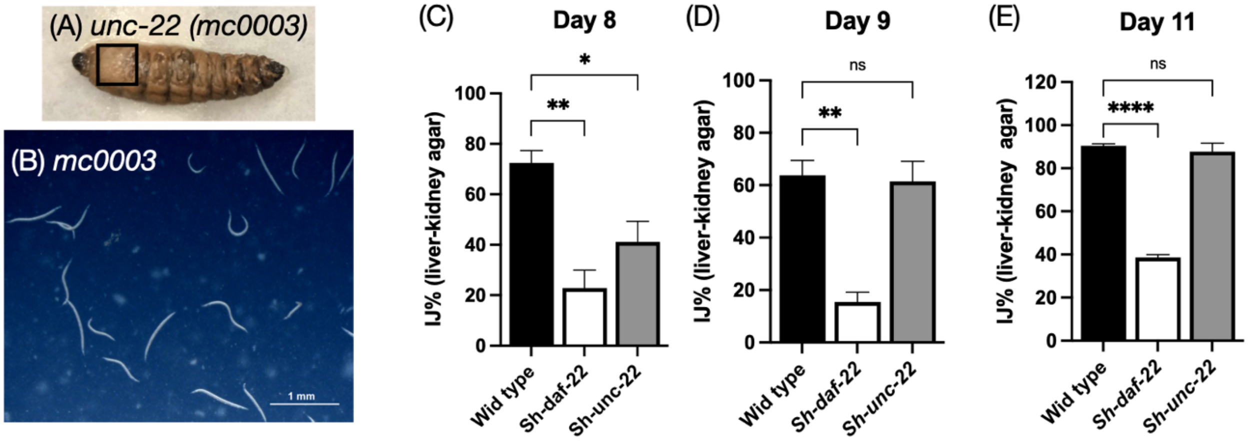


**Figure S4**: *In vivo* and *in vitro* IJ developmental defect is caused by *Sh-daf-22,* not *Sh-unc-22.* (A): *Sh-unc-22* mutant IJs emerged from insect cadaver and were resuspended in M9 buffer and examined by microscopy. (B): Emerged *Sh-unc-22* nematodes are primarily IJs. (C-E). Quantification of percentage of IJs in the nematode population on nutrient-rich liver kidney agar on 8 days (C), 9 days (D), and 11 days (E) of growth.
